## Supplementary material for "Damped oscillations of the probability of random events followed by absolute refractory period: exact analytical results": check_avp_formula_with_P_max_damping_coef.m

```
clear all;

T = 200; % [ms]
dt = 0.01; % [ms]
% tau_ref = 2; % [ms]
% p_s = 0.1;
tau_ref = 5; % [ms]
p_s = 0.01;

format long

N = floor(T/dt);
n_ref = floor(tau_ref/dt);
P = zeros(1,N);
Index = 1:N;
time = Index.*dt; % [ms]

P1 = @(p,k) p.*(1 - p).^(k-1);

P2 = @(p,k,n_ref,q) P1(p,k).*(1 + (k - n_ref - 1).*q);

P3 = @(p,k,n_ref,q) P1(p,k).*(1 + (k - n_ref - 1).*q + (1/2).*(k - 2.*n_ref - 2).*(k - 2.*n_ref - 1).*q.^2);

theor_stat_level = p_s/(1+n_ref*p_s)

for k=1:N
        
    if k<=(n_ref+1)
        P(k) = p_s*(1-p_s)^(k-1);  
    else
        P(k) = p_s*P(k-n_ref-1) + (1-p_s)*P(k-1);
    end;    
      
end;

P_max = zeros(1,N);

%========= 2nd interval over (n_ref +1) =================

q = p_s/(1-p_s)^(n_ref+1)

u = log(1/(1-p_s))

R = (q - u)/(q*u)

k_max2 = floor(n_ref + 1 + R)

P2_max = P2(p_s,k_max2,n_ref,q)

P2_max_check = ((p_s)^2/u)*(1-p_s)^(R-1)

P_max(k_max2) = P2_max;

D2 = (p_s/u)*(1-p_s)^(R-1)

D2_check = P2(p_s,n_ref + 1 + R,n_ref,q)/P1(p_s,1)

% ========= 3rd interval over (n_ref +1) ============

% a*k^2 + b*k + c = 0

a = 0.5*(q^2)*log(1-p_s)

b = q^2 - (2*n_ref + 3/2)*(q^2)*log(1-p_s) + q*log(1-p_s)

c = -(2*n_ref + 3/2)*(q^2) + (n_ref + 1)*(2*n_ref + 1)*(q^2)*log(1-p_s) + q - (n_ref + 1)*q*log(1-p_s) + log(1-p_s)

Det = b^2 - 4*a*c

k1 = (-b + sqrt(Det))/(2*a)

k2 = (-b - sqrt(Det))/(2*a)

k_max3 = floor(k2)

X = -1/2 + sqrt(1/4 + 1/u^2 -(2*n_ref + 1)/q - 1/q^2)

k_max3_check = 2*(n_ref+1) + R + X

P3_max = P3(p_s,k_max3,n_ref,q)

P3_max_check = (p_s*(1 - p_s)^(2*n_ref + 1 + R + X))*(1 + (n_ref + 1 + R + X)*q + (1/2)*(R + X)*(1 + R + X)*q^2)

P_max(k_max3) = P3_max;

D3 = P3(p_s,2*(n_ref+1) + R + X,n_ref,q)/P2(p_s,n_ref + 1 + R,n_ref,q)

D3_direct = (u/q^2)*(p_s*(1 - p_s)^X)*(1 + (n_ref + 1 + R + X)*q + (1/2)*(R + X)*(1 + R + X)*q^2)

D3_check = (p_s*(1 - p_s)^X)*(1/(u) + (X + 1/2))

D3_check2 = (p_s*(1 - p_s)^X)*(u/q)*(n_ref + 1 + 1/u + X + (1/2)*((R + X + 1/2)^2 - 1/4)*q)

OUT = [time; P];
fid = fopen('theor_event_probability_P_k_recur.txt','wt');
fprintf(fid,'%10.12f %10.12f\n',OUT); 
fclose(fid);

figure(1)
plot(time,P,'b',time,theor_stat_level.*ones(1,length(time)),'g',time,P_max,'r',time,P2_max.*ones(1,length(time)),'k',time,P3_max.*ones(1,length(time)),'k')
xlabel('t [ms]');
ylabel('Event probability P(t)');
```
