## Supplementary material for "Damped oscillations of the probability of random events followed by absolute refractory period: exact analytical results": check_avp_formula_explicite_form.m

```
% WARNING: At large time T, because of too large values of factorials the computation breaks, resulting in Inf/NaN output values.

T = 50; % [ms]
dt = 0.01; % [ms]
% tau_ref = 2; % [ms]
% p_s = 0.1;
tau_ref = 5; % [ms]
p_s = 0.01;

N = floor(T/dt);
n_ref = floor(tau_ref/dt);
P = zeros(1,N);
Index = 1:N;
time = Index.*dt; % [ms]

theor_stat_level = p_s/(1+n_ref*p_s);

P1 = @(p,k) p*(1 - p)^(k-1); % definition of function P1(p,k) = p*(1 - p)^(k-1)

q = p_s/(1-p_s)^(n_ref+1);

for k=1:N
    
    P(k) = P1(p_s,k);
    m = floor(k/n_ref);   
    S = 0;
    
    if (m>0)
        for i=1:m
            S = S + CC(k,i,n_ref)*q^(i);
            %S = S + CCC(k,i,n_ref)*q^(i); % the alternative with CCC function
        end;
        P(k) = P(k) + P1(p_s,k)*S;
    end;
    
    fprintf('%10.2f%%\n',100*k/N);
    
end;

OUT = [time; P];
fid = fopen('theor_event_probability_avp_P_k_explicit.txt','wt');
fprintf(fid,'%10.12f %10.12f\n',OUT); 
fclose(fid);

figure(1)
plot(time,P,'b',time,theor_stat_level.*ones(1,length(time)),'g')
xlabel('t [ms]');
ylabel('Event probability P(t)');
```
