## Supplementary material for "Damped oscillations of the probability of random events followed by absolute refractory period: exact analytical results": partial_formulas_for_P_k.m

```
T = 20; % [ms]
dt = 0.01; % [ms]
tau_ref = 2; % [ms]
p_s = 0.1;
% tau_ref = 5; % [ms]
% p_s = 0.01;

N = floor(T/dt);
n_ref = floor(tau_ref/dt);
P = zeros(1,N);
Index = 1:N;
time = Index.*dt; % [ms]

theor_stat_level = p_s/(1+n_ref*p_s);

P1 = @(p,k) p.*(1 - p).^(k-1);

P2 = @(p,k,n_ref) ((k - n_ref - 1).*(p.^2).*(1 - p).^(k - n_ref - 2) + P1(p,k));

P3 = @(p,k,n_ref) (0.5.*(k - 2.*n_ref - 2).*(k - 2.*n_ref - 1).*(p.^3).*(1 - p).^(k - 2.*n_ref - 3) + P2(p,k,n_ref));

for k=1:N
        
    if k<=(n_ref+1)
        P(k) = P1(p_s,k);  
    end;
    
    if ((n_ref+1)<=k)&&(k<=2*(n_ref+1))
        P(k) = P2(p_s,k,n_ref);  
    end;
    
    if (2*(n_ref+1)<=k)&&(k<=3*(n_ref+1))
        P(k) = P3(p_s,k,n_ref);  
    end;
      
end;

figure(1)
plot(time,P,'b',time,theor_stat_level.*ones(1,length(time)),'g')
xlabel('t [ms]');
ylabel('Event probability P(t)');

%=======================

k=1:(n_ref+1);
time = k.*dt;

PP = zeros(1,1);
PP(k) = P1(p_s,k);

figure(2)
plot(k.*dt,PP(k),'b')
xlabel('t = k*dt [ms]');
ylabel('P1_{k}');

OUT = [time; PP(k)];
fid = fopen('P1_k.txt','wt');
fprintf(fid,'%10.12f %10.12f\n',OUT); 
fclose(fid);

%=======================

k=(n_ref+1):(2*(n_ref+1));
time = k.*dt;

PP = zeros(1,1);
PP(k) = P2(p_s,k,n_ref);

figure(3)
plot(time,PP(k),'b')
xlabel('t = k*dt [ms]');
ylabel('P2_{k}');

OUT = [time; PP(k)];
fid = fopen('P2_k.txt','wt');
fprintf(fid,'%10.12f %10.12f\n',OUT); 
fclose(fid);

%=======================

k=(2*(n_ref+1)):(3*(n_ref+1));
time = k.*dt;

PP = zeros(1,1);
PP(k) = P3(p_s,k,n_ref);

figure(4)
plot(time,PP(k),'b')
xlabel('t = k*dt [ms]');
ylabel('P3_{k}');

OUT = [time; PP(k)];
fid = fopen('P3_k.txt','wt');
fprintf(fid,'%10.12f %10.12f\n',OUT); 
fclose(fid);
```
