## Supplementary material for "Damped oscillations of the probability of random events followed by absolute refractory period: exact analytical results": event_probability_simulation_with_raster.m

```
T = 100; % simulation time [ms]
dt = 0.01; % time step [ms]
% tau_ref = 2; % absolute refractory period [ms]
% p_s = 0.1; % elementary (or unconditional) probability of an event occurrence during dt
tau_ref = 5; % [ms]
p_s = 0.01;

N = floor(T/dt);
Index = 1:N;
time = Index.*dt; % [ms]

Stat = zeros(1,N);

NN = 10000; % number of independent neurons
t_sp = -tau_ref*ones(1,NN); % initial condition

fid = fopen('raster_of_events.txt','wt');

for i=1:N

t = i*dt;
    
for j=1:NN    
    if (rand(1)<=p_s)&&(t>(t_sp(j)+tau_ref))
        t_sp(j) = t;
        Stat(i) = Stat(i) + 1;
        fprintf(fid,'%10.12f %d\n',t,j); % raster of events
    end;
end;

fprintf('%10.2f%%\n',100*i/N);

end;

fclose(fid);

Prob = Stat./NN;

OUT = [time; Prob];
fid = fopen('event_probability_simulation_with_raster.txt','wt');
fprintf(fid,'%10.12f %10.12f\n',OUT); 
fclose(fid);

figure(1)
plot(time,Prob,'b',time,mean(Prob).*ones(1,length(time)),'g')
xlabel('t [ms]');
ylabel('Event probability P(t)');
```
