## Supplementary material for "Damped oscillations of the probability of random events followed by absolute refractory period: exact analytical results": check_avp_formula_via_sums_diff.m

```
T = 200; % [ms]
dt = 0.01; % [ms]
% tau_ref = 2; % [ms]
% p_s = 0.1;
tau_ref = 5; % [ms]
p_s = 0.01;

N = floor(T/dt);
n_ref = floor(tau_ref/dt);
P = zeros(1,N);
Index = 1:N;
time = Index.*dt; % [ms]

S = 0;
SS = 0;

theor_stat_level = p_s/(1+n_ref*p_s)

for k=1:N
    
    S = 0;
    if k>1
        for j=1:k-1
            S = S + P(j);
        end;
    else
        S = 0;
    end;
        
    SS = 0;
    if k>(n_ref+1)
        for j=1:(k-n_ref-1)
            SS = SS + P(j);
        end;
    else
        SS = 0;
    end;
        
    P(k) = p_s*(1-(S-SS));     
      
end;

OUT = [time; P];
fid = fopen('theor_event_probability_P_k_via_sums_diff.txt','wt');
fprintf(fid,'%10.12f %10.12f\n',OUT); 
fclose(fid);

figure(1)
plot(time,P,'b',time,theor_stat_level.*ones(1,length(time)),'g')
xlabel('t [ms]');
ylabel('Event probability P(t)');
```
