## Supplementary material for "Damped oscillations of the probability of random events followed by absolute refractory period: exact analytical results": renewal_density_analyt_formula.m

```
clear all;

T = 100; % [ms]
dt = 0.01; % [ms]
% tau_ref = 2; % [ms]
% p_s = 0.1;
tau_ref = 5; % [ms]
p_s = 0.01;

N = floor(T/dt);
n_ref = floor(tau_ref/dt);
P = zeros(1,N);
Index = 1:N;
time = Index.*dt; % [ms]

theor_stat_level = p_s/(1+n_ref*p_s);

for k=1:N
        
    if k<=(n_ref+1)
        P(k) = p_s*(1-p_s)^(k-1);  
    else
        P(k) = p_s*P(k-n_ref-1) + (1-p_s)*P(k-1);
    end;    
      
end;

OUT = [time; P];
fid = fopen('theor_firing_probability_P_k_recur.txt','wt');
fprintf(fid,'%10.12f %10.12f\n',OUT); 
fclose(fid);

figure(1)
plot(time,P,'b',time,theor_stat_level.*ones(1,length(time)),'g')
xlabel('t [ms]');
ylabel('Event probability P(t)');

% ========= Computing the probability P_k through renewal density h(t) for Poisson + dead time tau_ref =========

mu = p_s/dt; % parameter in the Poisson distribution
n_max = 1000; % number of terms in the sum of f_n over n

h = zeros(1,N); % renewal density
P_rd = zeros(1,N); % probability P_k determined through renewal density

for k=1:N

t = k*dt;

for n=1:n_max
S = 0;
if t>=n*tau_ref
    S = mu*(((mu*(t-n*tau_ref))^(n-1))/factorial(n-1))*exp(-mu*(t-n*tau_ref));
end;
h(k) = h(k) + S;
end; 

h(k) = h(k);

P_rd(k)=h(k)*dt;

fprintf('%10.2f%%\n',100*k/N);

end;

figure(2)
plot(time,P_rd,'k',time,theor_stat_level.*ones(1,length(time)),'g')
xlabel('t [ms]');
ylabel('P_{rd}(t)');

figure(3)
plot(time,P,'b',time,P_rd,'k',time,theor_stat_level.*ones(1,length(time)),'g')
xlabel('t [ms]');
ylabel('P(t), P_{rd}(t)');

OUT = [time; P_rd];
fid = fopen('P_rd__renewal_density_mult_by_dt.txt','wt');
fprintf(fid,'%10.12f %10.12f\n',OUT); 
fclose(fid);
```
