## Supplementary material for "Damped oscillations of the probability of random events followed by absolute refractory period: exact analytical results": Supplementary_Material__Derivation_of_the_damping_coefficients.pdf

**Supplementary Material: Derivation of the damping coefficients for the 2nd and 3rd peaks of the event probability**

**1. Prerequisites.** The probability of spike generation in the interval $1 \le k \le n_{ref}+1$ is given by

$$P_k^{(1)} = p_s(1-p_s)^{k-1}.$$

Next, at $n_{ref}+1 \le k \le 2(n_{ref}+1)$ for the probability one gets

$$P_k^{(2)} = (k-n_{ref}-1)p_s^2(1-p_s)^{k-n_{ref}-2} + P_k^{(1)} = P_k^{(1)}[1+(k-n_{ref}-1)q],$$

where $q = p_s(1-p_s)^{-(n_{ref}+1)}$. Finally, at $2(n_{ref}+1) \le k \le 3(n_{ref}+1)$ one has

$$P_k^{(3)} = \frac{1}{2}(k-2n_{ref}-2)(k-2n_{ref}-1)p_s^3(1-p_s)^{k-2n_{ref}-3} + P_k^{(2)} =$$
$$= P_k^{(1)}[1+(k-n_{ref}-1)q + \frac{1}{2}(k-2n_{ref}-2)(k-2n_{ref}-1)q^2].$$

These explicit formulas allow analytical calculation of the relative damping of the 2nd and 3rd peaks of $P_k$. The damping is defined as the ratio of amplitudes of the adjacent decreasing peaks,

$$D_{i+1} = P_{\max}^{(i+1)}/P_{\max}^{(i)},$$

where $P_{\max}^{(i)} \equiv P_{k_{\max i}}^{(i)}$ and $k_{\max i}$ is the position of the $i$-th peak.

**2. Derivation.** The position of the 1st peak obviously corresponds to $k_{\max 1} = 1$ leading to $P_{\max}^{(1)} = p_s$. Further, assuming that $k$ is a continuous variable and calculating $dP_k^{(2)}/dk = 0$, for $n_{ref}+1 \le k \le 2(n_{ref}+1)$ one gets a linear algebraic equation for the position $k_{\max 2}$ of the 2nd peak. In particular, starting with

$$P_k^{(2)} = (k-n_{ref}-1)p_s^2(1-p_s)^{k-n_{ref}-2} + p_s(1-p_s)^{k-1}$$

and using formula

$$\frac{d}{dx}a^{f(x)} = a^{f(x)}\ln(a)\frac{df}{dx},$$

one gets

$$\frac{dP_k^{(2)}}{dk} = p_s^2(1-p_s)^{k-n_{ref}-2} + (k-n_{ref}-1)p_s^2(1-p_s)^{k-n_{ref}-2}\ln(1-p_s) + p_s(1-p_s)^{k-1}\ln(1-p_s) =$$
$$= p_s(1-p_s)^{k-1}\left[p_s(1-p_s)^{-(n_{ref}+1)} + (k-n_{ref}-1)p_s(1-p_s)^{-(n_{ref}+1)}\ln(1-p_s) + \ln(1-p_s)\right] = 0$$

Denoting $q = p_s(1-p_s)^{-(n_{ref}+1)}$ and $u = \ln[1/(1-p_s)]$, we arrive at linear algebraic equation for $k$

$$q - (k-n_{ref}-1)qu - u = 0,$$

Its solution is straightforward,

$$k \equiv k_{\max 2} = n_{ref} + 1 + 1/u - 1/q = n_{ref} + 1 + R,$$

where $R = 1/u - 1/q$. Accordingly, the amplitude of the 2nd peak is given by

$$P_{\max}^{(2)} \equiv P_{k_{\max 2}}^{(2)} = Rp_s^2(1-p_s)^{R-1} + p_s(1-p_s)^{n_{ref}+R}.$$

Expressing

$$(1-p_s)^{n_{ref}} = \frac{p_s}{q(1-p_s)},$$

one finally gets

$$P_{\max}^{(2)} = Rp_s^2(1-p_s)^{R-1} + \frac{p_s^2}{q}(1-p_s)^{R-1} = (R+\frac{1}{q})p_s^2(1-p_s)^{R-1} = \frac{p_s^2}{u}(1-p_s)^{R-1}.$$

Given that $P_{\max}^{(1)} = p_s$, for the relative damping of the second peak we get

$$D_2 = P_{\max}^{(2)}/P_{\max}^{(1)} = \frac{p_s}{u}(1-p_s)^{R-1}.$$

Turning to the 3rd peak, for $2(n_{ref}+1) \leq k \leq 3(n_{ref}+1)$ we have

$$P_k^{(3)} = \frac{1}{2}(k-2n_{ref}-2)(k-2n_{ref}-1)p_s^3(1-p_s)^{k-2n_{ref}-3} + P_k^{(2)}.$$

Let us calculate $dP_k^{(3)}/dk = 0$:

$$\frac{dP_k^{(3)}}{dk} = \frac{1}{2}(2k-4n_{ref}-3)p_s^3(1-p_s)^{k-2n_{ref}-3} +$$
$$+\frac{1}{2}(k-2n_{ref}-2)(k-2n_{ref}-1)p_s^3(1-p_s)^{k-2n_{ref}-3}\ln(1-p_s) + \frac{dP_k^{(2)}}{dk},$$
$$\frac{dP_k^{(3)}}{dk} = p_s(1-p_s)^{k-1}[(2k-4n_{ref}-3)-(k-2n_{ref}-2)(k-2n_{ref}-1)u]\frac{1}{2}q^2 + \frac{dP_k^{(2)}}{dk} = 0.$$

Substituting here the above expression for $dP_k^{(2)}/dk$, now we get quadratic algebraic equation for $k$,

$$[(2k-4n_{ref}-3)-(k^2-4n_{ref}k+4n_{ref}^2-3k+6n_{ref}+2)u]\frac{1}{2}q^2 + q - (k-n_{ref}-1)qu - u = 0,$$
$$k^2(uq^2) - 2k(q^2+(2n_{ref}+\frac{3}{2})uq^2-qu) + 2[(2n_{ref}+\frac{3}{2})q^2+(2n_{ref}^2+3n_{ref}+1)uq^2-q-(n_{ref}+1)qu+u] = 0.$$

Denoting $x = \frac{1}{2}q^2u$, $y = q^2 + (2n_{ref}+\frac{3}{2})q^2u - qu$, and

$$z = -(2n_{ref}+\frac{3}{2})q^2 - (2n_{ref}^2+3n_{ref}+1)uq^2 + q + (n_{ref}+1)qu - u =$$
$$= -(2n_{ref}+\frac{3}{2})q^2 - (n_{ref}+1)(2n_{ref}+1)q^2u + q + (n_{ref}+1)qu - u,$$

we arrive at

$$xk^2 - yk - z = 0,$$

the roots of which are

$$k = \frac{y \pm \sqrt{y^2+4xz}}{2x}.$$

Taking the positive root, we get

$$k_{\max 3} = \frac{y+\sqrt{y^2+4xz}}{2x} = \frac{y}{2x} + \sqrt{\left(\frac{y}{2x}\right)^2 + \frac{z}{x}},$$

where

$$\frac{y}{2x} = \frac{q^2+(2n_{ref}+\frac{3}{2})q^2u-qu}{q^2u} = \frac{1}{u} + 2n_{ref} + \frac{3}{2} - \frac{1}{q} = R + 2n_{ref} + \frac{3}{2} = R + 2(n_{ref}+1) - \frac{1}{2},$$

$$\frac{z}{x} = \frac{-(2n_{ref}+\frac{3}{2})q^2 - (n_{ref}+1)(2n_{ref}+1)q^2u + q + (n_{ref}+1)qu - u}{\frac{1}{2}q^2u} =$$
$$= -\frac{2(2n_{ref}+1)}{u} - \frac{1}{u} - 2(n_{ref}+1)(2n_{ref}+1) + \frac{2}{qu} + \frac{2(n_{ref}+1)}{q} - \frac{2}{q^2}$$

$$\left(\frac{y}{2x}\right)^2 = \left(\frac{1}{u} + (2n_{ref}+1) + \frac{1}{2} - \frac{1}{q}\right)^2 =$$
$$= \frac{1}{u^2} + (2n_{ref}+1)^2 + \frac{1}{4} + \frac{1}{q^2} + 2(2n_{ref}+1)\left(\frac{1}{u} - \frac{1}{q}\right) + (2n_{ref}+1) + \frac{1}{u} - \frac{1}{q} - \frac{2}{qu},$$

$$\left(\frac{y}{2x}\right)^2 + \frac{z}{x} = \frac{1}{u^2} + (2n_{ref}+1)^2 + \frac{1}{4} + \frac{1}{q^2} + 2(2n_{ref}+1)\left(\frac{1}{u} - \frac{1}{q}\right) + (2n_{ref}+1) + \frac{1}{u} - \frac{1}{q} - \frac{2}{qu} -$$
$$- \frac{2(2n_{ref}+1)}{u} - \frac{1}{u} - 2(n_{ref}+1)(2n_{ref}+1) + \frac{2}{qu} + \frac{2(n_{ref}+1)}{q} - \frac{2}{q^2}$$
$$= \frac{1}{u^2} + \frac{1}{4} + 2(2n_{ref}+1)\left(\frac{1}{u} - \frac{1}{q}\right) - \frac{1}{q} - \frac{2(2n_{ref}+1)}{u} + \frac{2(n_{ref}+1)}{q} - \frac{1}{q^2} =$$
$$= \frac{1}{u^2} + \frac{1}{4} - \frac{1}{q^2} + \frac{4n_{ref}+2-4n_{ref}-2}{u} - \frac{4n_{ref}+2+1-2n_{ref}-2}{q} =$$
$$= \frac{1}{u^2} + \frac{1}{4} - \frac{1}{q^2} - \frac{(2n_{ref}+1)}{q}.$$

Denoting

$$X = -\frac{1}{2} + \sqrt{\left(\frac{y}{2x}\right)^2 + \frac{z}{x}} = -\frac{1}{2} + \sqrt{\frac{1}{4} + \frac{1}{u^2} - \frac{1}{q^2} - \frac{(2n_{ref}+1)}{q}},$$

we finally get

$$k_{\max 3} = R + 2(n_{ref}+1) + X.$$

Substituting $k_{\max 3}$ into the expression for $P_k^{(3)}$ yields the amplitude of the 3rd peak,

$$P^{(3)}_{k_{\max 3}} = P^{(1)}_{k_{\max 3}}[1 + (k_{\max 3} - n_{ref} - 1)q + \frac{1}{2}(k_{\max 3} - 2n_{ref} - 2)(k_{\max 3} - 2n_{ref} - 1)q^2] =$$
$$= P^{(1)}_{k_{\max 3}}[1 + (n_{ref} + 1 + R + X)q + \frac{1}{2}(R+X)(1+R+X)q^2],$$

where $P^{(1)}_{k_{\max 3}} = p_s(1-p_s)^{k_{\max 3}-1} = p_s(1-p_s)^{R+X+2n_{ref}+1}$

The damping coefficient of the 3rd peak,

$$D_3 = P^{(3)}_{k_{\max 3}}/P^{(2)}_{k_{\max 2}} = \frac{p_s(1-p_s)^{R+X+2n_{ref}+1}[1 + (n_{ref}+1+R+X)q + \frac{1}{2}(R+X)(1+R+X)q^2]}{\frac{p_s^2}{u}(1-p_s)^{R-1}} =$$
$$= \frac{u}{p_s}(1-p_s)^{X+2(n_{ref}+1)}[1 + (n_{ref}+1+R+X)q + \frac{1}{2}(R+X)(1+R+X)q^2] =$$
$$= p_s(1-p_s)^X \frac{u}{q^2}[1 + (n_{ref}+1+R+X)q + \frac{1}{2}(R+X)(1+R+X)q^2].$$

To get the last formula, we have used the definition of $q = p_s(1-p_s)^{-(n_{ref}+1)}$.

The cumbersome expression in $D_3$

$$\frac{u}{q^2}[1 + (n_{ref}+1+R+X)q + \frac{1}{2}(R+X)(1+R+X)q^2] \equiv Q$$

can be further transformed into a compact form taking into account that

$$X^2 = \left(-\frac{1}{2} + \sqrt{\frac{1}{4} + \frac{1}{u^2} - \frac{1}{q^2} - \frac{(2n_{ref}+1)}{q}}\right)^2 = \frac{1}{u^2} - \frac{1}{q^2} - \frac{(2n_{ref}+1)}{q} - X,$$

and using the definition for $R = 1/u - 1/q$.

Indeed,

$$
\begin{aligned}
Q &= [\frac{u}{q^2} + (n_{ref} + 1 + \frac{1}{u} - \frac{1}{q} + X)\frac{u}{q} + \frac{1}{2}(\frac{1}{u} - \frac{1}{q} + X)(1 + \frac{1}{u} - \frac{1}{q} + X)u] = \\
&= (n_{ref} + 1)\frac{u}{q} + \frac{1}{q} + X\frac{u}{q} + \frac{1}{2}(1 - \frac{u}{q} + uX)(1 + \frac{1}{u} - \frac{1}{q} + X).
\end{aligned}
$$

Expanding the parentheses in the last formula, we get

$$
\begin{aligned}
(1 - \frac{u}{q} + uX)(1 + \frac{1}{u} - \frac{1}{q} + X) &= 1 - \frac{u}{q} + uX + \frac{1}{u}(1 - \frac{u}{q} + uX) - \frac{1}{q}(1 - \frac{u}{q} + uX) + X(1 - \frac{u}{q} + uX) = \\
&= 1 - \frac{u}{q} + uX + \frac{1}{u} - \frac{1}{q} + X - \frac{1}{q} + \frac{u}{q^2} - \frac{u}{q}X + X - X\frac{u}{q} + uX^2 = \\
&= 1 - \frac{u}{q} + uX + \frac{1}{u} - \frac{2}{q} + 2X + \frac{u}{q^2} - 2\frac{u}{q}X + uX^2 = \\
&= 1 - \frac{u}{q} + uX + \frac{1}{u} - \frac{2}{q} + 2X + \frac{u}{q^2} - 2\frac{u}{q}X + u\left(\frac{1}{u^2} - \frac{1}{q^2} - \frac{(2n_{ref} + 1)}{q} - X\right) = \\
&= 1 - \frac{u}{q} + uX + \frac{1}{u} - \frac{2}{q} + 2X + \frac{u}{q^2} - 2\frac{u}{q}X + \frac{1}{u} - \frac{u}{q^2} - 2n_{ref}\frac{u}{q} - \frac{u}{q} - uX = \\
&= 1 - 2(n_{ref} + 1)\frac{u}{q} + \frac{2}{u} - \frac{2}{q} + 2X - 2\frac{u}{q}X
\end{aligned}
$$

Turning back to $Q$,

$$
Q = (n_{ref} + 1)\frac{u}{q} + \frac{1}{q} + X\frac{u}{q} + \frac{1}{2} - (n_{ref} + 1)\frac{u}{q} + \frac{1}{u} - \frac{1}{q} + X - \frac{u}{q}X = \frac{1}{2} + \frac{1}{u} + X.
$$

Thus, the final expression for the damping coefficient of the 3rd peak is given by

$$
D_3 = P^{(3)}_{k_{\max 3}} / P^{(2)}_{k_{\max 2}} = Qp_s(1 - p_s)^X = (\frac{1}{2} + \frac{1}{u} + X)p_s(1 - p_s)^X.
$$
